# Reference Gene Validation in Energy Cane Under Smut Infection and Insights into Antioxidant Gene Expression

**DOI:** 10.1101/2025.10.24.683138

**Authors:** G.S. Crestana, M. Ferreira, J.D. Ferreti, J.A. Bressiani, S. Creste, C.B. Monteiro-Vitorello

## Abstract

*Sporisorium scitamineum* is the causal agent of smut disease affecting sugarcane and energy cane crop production. Despite the frequent use of traditional reference genes for gene expression normalization by RT-qPCR, their stability can vary across genotypes and stress conditions. Here, we evaluated the stability of seven commonly used reference genes (*GAPDH*, *SAMDC*, *UBQ*, *25S rRNA*, *αTUB*, *SARMp1*, and *ACAD*) and two additional candidates (*EF5A-1* and *SDH1-1*) identified from RNA-Seq data in two contrasting energy cane genotypes (Vertix 1, smut-susceptible; Vertix 2, smut-resistant). Stability was assessed at three infection time points (12 hours post-inoculation, 48 hpi, and 5 days post-inoculation) separately and also on a combined dataset comprising all time points, under mock and smut-inoculated conditions. Gene stability, evaluated through multiple approaches (coefficient of variation, RefFinder, GeNorm) and consolidated using RankAggreg, revealed genotype and condition-dependent variability, with no universal reference gene set for most conditions. *GAPDH*, commonly used in sugarcane studies, showed the highest variation in our experiments, reinforcing the need for genotype and condition-specific validation. The best-ranked reference genes were used to quantify ROS-related defense responses. Vertix 2 (Vx2) exhibited coordinated induction of *SOD*, *POX5*, and *CATB*, whereas Vertix 1 (Vx1) showed delayed or reduced activation of these genes alongside a compensatory up-regulation of *GSTt3*. By validating reliable endogenous genes from literature and RNA-Seq data, this study ensures accurate gene expression quantification under biotic stress and highlights antioxidant mechanisms as contributors to smut resistance in *S. scitamineum*–energy cane interactions.

## Introduction

Sugarcane breeding programs have developed cultivars for high sucrose content in the culms used primarily for sugar and first-generation (1G) ethanol production. However, because of the crop’s significant potential for bioenergy, breeding efforts have led to developing novel genotypes for producing cellulosic or second-generation (2G) ethanol, such as energy cane [Cursi et al., 2022; de Abreu et al., 2020]. These genotypes have a genetic background originating from the first generation of crosses between modern sugarcane varieties and *Saccharum spontaneum*, whose larger genetic contribution results in the higher biomass production observed in energy canes. By comparison, the modern sugarcane cultivars are predominantly *S. officinarum* (2n = 80; ∼90%) hybrids with limited *S. spontaneum* introgression (2n = 40–128; ∼10%), and contributions from *S. barberi* and *S. sinense* [Amalraj & Balasundaram, 2006; Nair, 2012; Healey et al., 2024; Thirugnanasambandam, Hoang & Henry, 2018; Zhang et al., 2018]. Although energy cane genotypes have significant potential for biomass production, they show moderate to high susceptibility to smut disease [Diniz et al., 2019]. Smut is widespread across sugarcane-growing regions worldwide, and genetic resistance remains the primary management strategy [Bhuiyan et al., 2021; Sundar et al., 2012].

In resistant sugarcane genotypes, smut colonization does not lead to whip formation, the fungal reproductive structure and characteristic symptom of the disease, resulting in no crop losses [Rajput et al. 2022]. Upon pathogen perception, resistant genotypes trigger an intense oxidative burst at the infection site with H_2_O_2_ accumulation at three time points related to the three phases of fungal development (6, 48, and 72 hpi), a response not observed in susceptible plants and known to signal defense response [Peters et al., 2017; Peters et al., 2020]. Transcriptomic data produced using genotypes with different genetic backgrounds reinforce this typical response upon smut infection [Agisha et al., 2022]. Some responsive genes encode Respiratory burst oxidase homolog (*RBHO*), Superoxide dismutase (*SOD*), Catalase (*CAT*), Glutathione S-transferase (*GSTt3*), Thioredoxin (*TRX*), and Peroxidase 5 (*POX5*), indicating the activation of the enzymatic antioxidant system and peroxide production. Different genotypes of modern cultivars and *S. spontaneum* infected with smut have similar responses varying among resistant and susceptible plants regarding the timing of the events. Resistant genotypes trigger key responses earlier and more consistently than susceptible ones, which often display delayed or uncoordinated activation [Hu et al., 2024]. So far, cane energy’s responses to smut are poorly understood.

Among the applications of Reverse Transcription quantitative PCR (RT-qPCR), measuring and comparing gene expression levels by quantifying RNA transcripts offers a highly sensitive and specific method, often used to validate RNA-seq data [Borrás-Hidalgo et al., 2005; Peters et al., 2017; McNeil et al., 2018]. The reference genes encoding *GAPDH*, *ACT*, *EF1α*, *UBQ*, and *25S ribosomal RNA* are references for sugarcane gene expression studies, including those analyzing the responses to smut infection [de Andrade et al., 2017; Guo et al., 2014; Hu et al., 2024; Huang et al., 2018]. However, gene expression may vary depending on experimental conditions, stress, and pathogens’ development patterns interacting with plants. Considering sugarcane genomes’ highly polyploid and aneuploid nature, selecting reference genes is more crucial for expression data analysis. Thus, even reference genes commonly used in related systems require validation under specific experimental conditions. Nevertheless, no validated set of reference genes is available for studying the energy cane–smut interaction. Regarding the statistical approaches commonly used to assess reference gene stability, several algorithms are available, including *geNorm* [Vandesompele et al., 2002], *NormFinder* [Andersen et al., 2004], *BestKeeper* [Pfaffl et al., 2004], the delta-Ct method [Silver et al., 2006], and *RefFinder* [Xie et al., 2023].

This work focused on identifying and validating suitable reference genes to analyze experimental data of the interaction between energy cane and *Sporisorium scitamineum*. Two contrasting energy cane genotypes (smut-resistant and smut-susceptible) were inoculated with smut and sampled across multiple time points during infection. Candidate reference genes selected among those used in sugarcane and from our RNA-Seq data were analyzed using various statistical algorithms. The most stable genes were applied to validate the expression of Reactive oxygen species (ROS)-related defense genes in energy cane, providing a proof of concept and insights into the early molecular responses involved in resistance.

## Material and Methods

### Ethics statement

Healthy energy cane buds used to conduct the experiments were gently provided by the Agronomic Institute of Campinas (IAC), Ribeirão Preto, São Paulo, Brazil, in collaboration with Nufarm Brasil. SSC04 diploid teliospores previously obtained [Benevenuto et al., 2016] were stored and maintained in the Genetics Department of ESALQ/USP (Genomics Group Laboratory). No special permits were necessary for the use of teliospores or cane collection.

### Experimental design and Biological material

We used single-bud setts of energy cane genotypes Vertix 1 (smut-susceptible) and Vertix 2 (smut-resistant). Buds were subjected to surface-disinfected by thermal (52 °C, 30 min) and chemical treatment (4% sodium hypochlorite, 10 min), arranged in trays containing vermiculite (2 cm depth) and incubated 24 hours in a humid chamber at 28 °C for pre-germination. Teliospores of *S. scitamineum* (strain SSC04) were previously obtained from diseased plants of RB925345 genotype, and viability was verified, with germination rates exceeding 90%. Inoculation was performed by applying a 10 µL drop of spore suspension containing 1 × 10^9^ spores/mL in 0.85% NaCl and 0.01% (v/v) Tween-20 to the bud base after causing a needle injury. Mock controls received 0.85% NaCl and 0.01% (v/v) Tween-20 only. Inoculated bud sets were maintained at 28 °C under conditions of 12 h light/12 h dark and 85% relative humidity. Three independent biological replicates (pools of 10 buds) were collected at 12 hpi, 48 hpi, and 5 dpi and stored at −80 °C. Time points were selected based on ROS production in bud scales of sugarcane, as described by Peters et al. (2017).

### Selection of candidate reference genes using publicly available RNA-seq data

Candidate reference primers commonly used in sugarcane and in studies addressing the interaction with *S. scitamineum* were initially selected from the literature. To further assess their stability and also identify new candidate reference genes for this study, publicly available RNA-seq data (FASTQ format) from 12 samples of the energy cane genotypes Vx1 and Vx2 collected 48 hours post inoculation with *S. scitamineum* were obtained from the NCBI SRA database (BioProject PRJNA1310642; BioSamples SAMN50776885–SAMN50776896; SRA runs SRR35122924–SRR35122927). Each genotype included three biological replicates under Mock and inoculated conditions. The RNA-seq data correspond to libraries constructed using the Illumina TruSeq Stranded mRNA Library Prep Kit and sequenced on the Illumina platform, generating paired-end reads (2×100 bp).

Read quality was assessed using *FastQC* v0.11.5 [Simon, A., 2010], and adapter trimming as well as removal of low-quality reads (average Q < 20), reads containing uncalled bases (N), and sequences shorter than 20 bp were performed using *Cutadapt* v1.18 [Martin, 2011]. We then aligned reads to the *Saccharum* hybrid cultivar R570 reference genome assembly [Healey et al., 2024] using *Hisat2* v2.1.0 [Kim & Salzberg, 2015] with default parameters. Gene-level read counts were generated with *FeatureCounts* v1.6.0 from the *Subread* package [Liao et al., 2014] using the corresponding R570 gene annotation file and the “-p” option for paired-end reads. Normalization of read counts was carried out using the *EdgeR* v4.6.3 package [Robinson et al., 2010] within the R/Bioconductor environment v4.5.1, based on counts per million (CPM) values normalized by the trimmed mean of M-values (TMM) method.

Expression stability was assessed across all treatments and genotypes. For each gene, we calculated the coefficient of variation (CV) as the ratio of the standard deviation to the mean of normalized read counts across all biological replicates. Genes showing CV ≤ 10% were considered stable and retained as candidates for subsequent validation by RT-qPCR.

### *In silico* analysis and primer design

Reference genes selected for RT-qPCR stability analysis included candidates previously reported as suitable for gene expression normalization in sugarcane, along with additional genes identified from RNA-Seq data (Table 1). Primer sequences for *GAPDH* (glyceraldehyde-3-phosphate dehydrogenase), *SAMDC* (S-adenosylmethionine decarboxylase), *UBQ* (ubiquitin), *αTUB* (alpha-tubulin), and *25S rRNA* (25S ribosomal RNA) were retrieved from the corresponding literature, whereas coding sequences (CDS) of *SDH1-1* (succinate dehydrogenase 1-1) and *EF5A-1* (elongation factor 5A-1) were obtained from the reference genome of the modern sugarcane cultivar R570 [Healey et al., 2024], available at the Sugarcane Genome Hub (https://sugarcane-genome.cirad.fr/). Homologous sequences were identified for *SDH1-1* and *EF5A-1* using BLASTn with an E-value cutoff of 1e–05 (parameter: -outfmt “6 std qcovs”). The BLAST hits were further filtered based on expression in both genotypes and a coefficient of variation (CV) ≤ 10%, then grouped into gene-specific sequence sets for multiple sequence alignment (MSA). Nucleotide sequences were aligned using ClustalW with CLC Genomics Workbench v 20.0.4 (QIAGEN, default parameters), and conserved regions shared among homologs were used for primer design. Primers were designed and evaluated in NetPrimer (www.premierbiosoft.com/netprimer) with the following parameters: 58–62 °C melting temperature (Tm), 18–22 bp length, 100–200 bp amplicon size, and optimized GC content, stability, and minimal primer-dimer interactions.

**Table 1.**
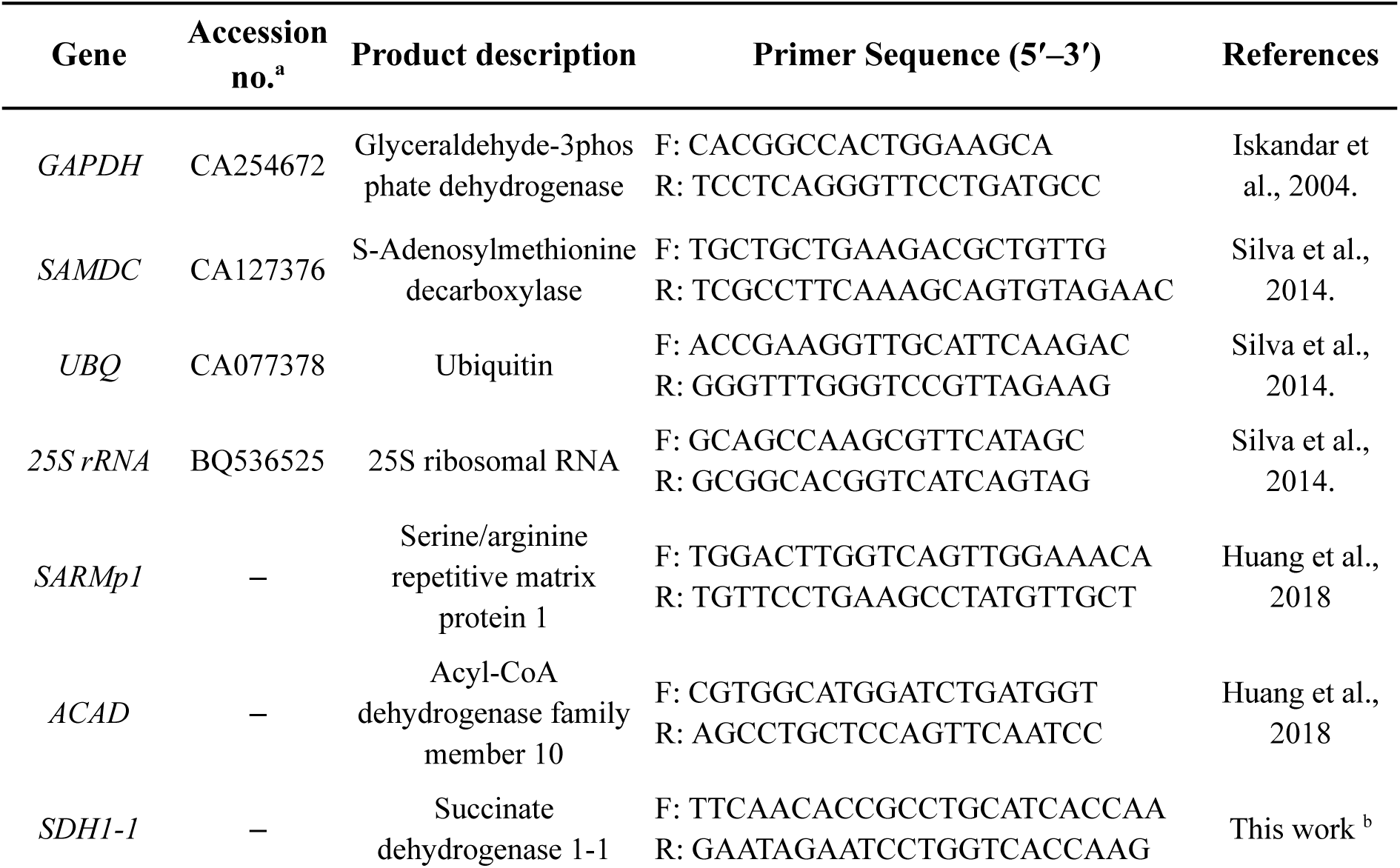
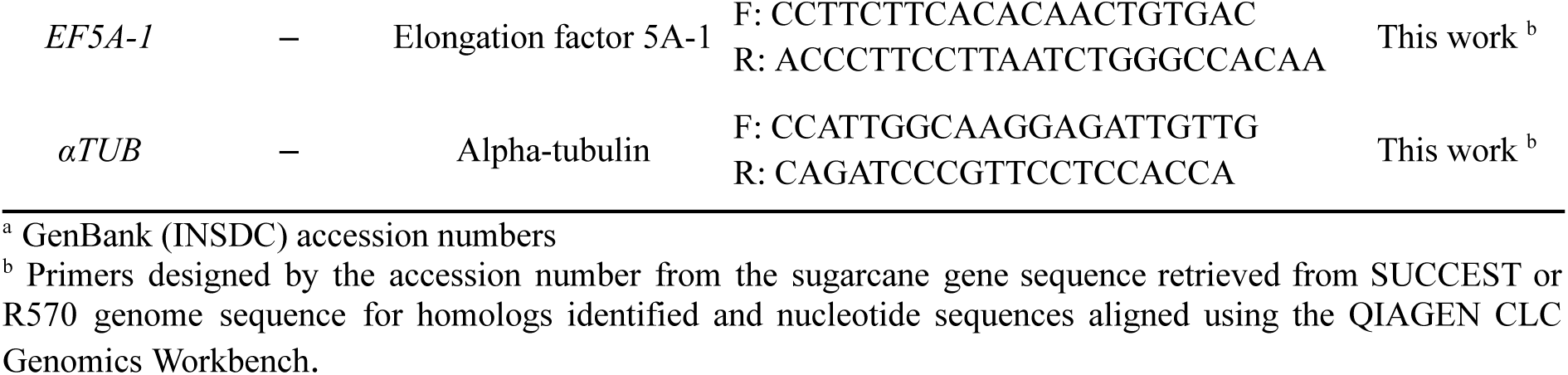
Reference Genes: gene name, accession number, product description, primer sequences, and literature reference.

Genes encoding antioxidant enzymes involved in of ROS metabolism were selected based on previous studies reporting differential expression from RNAseq data of sugarcane plants infected with *S. scitamineum* [Schaker et al., 2016] and on proteomics analysis in which differences in protein abundance were observed [Peters et al., 2017]. As described by Peters et al. (2017), the primer sequences (Table 2) were manually designed, and their quality was verified using the software Gene Runner (http://www.generunner.net/) and Beacon DesignerTM Free Edition (http://www.premierbiosoft.com).

**Table 2.**
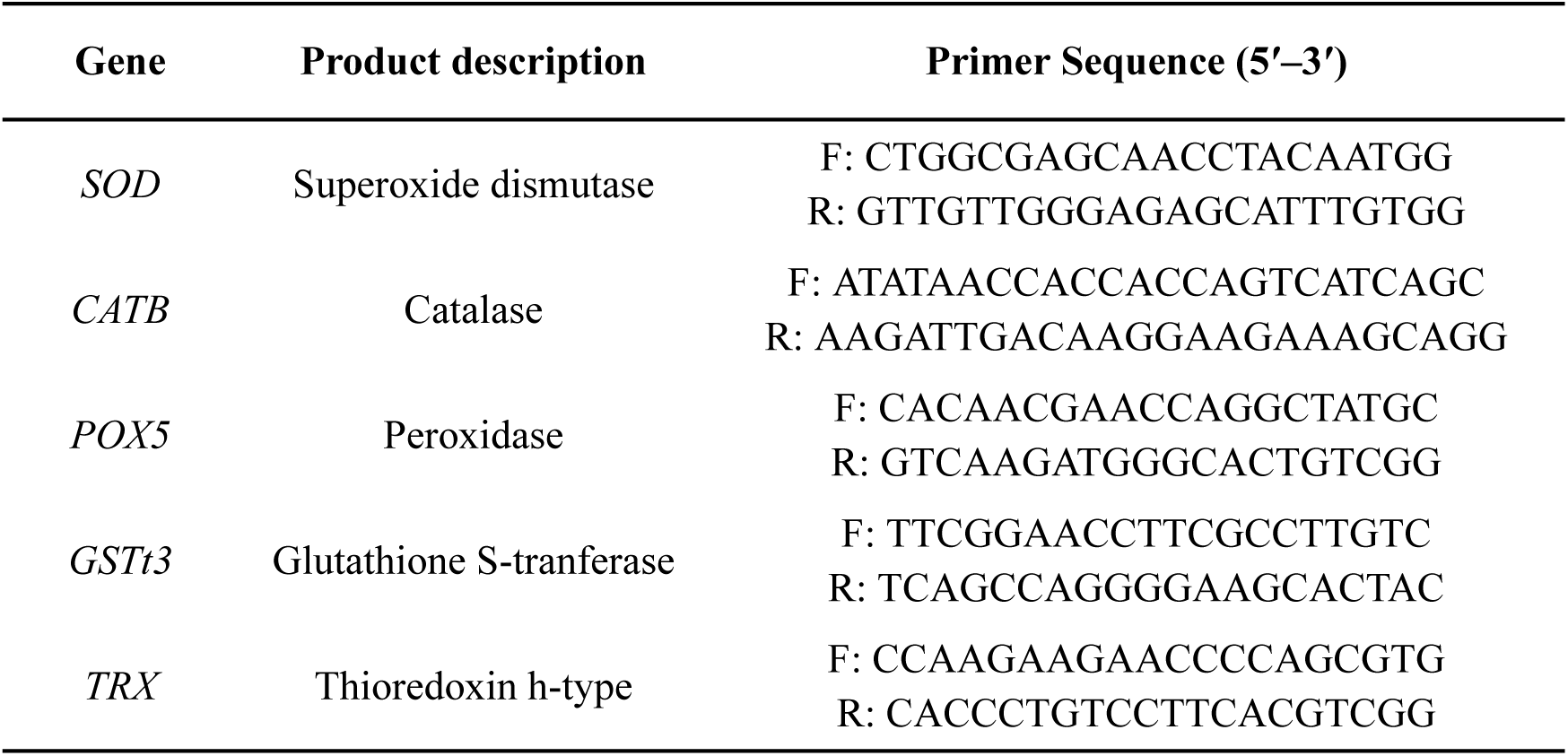
Antioxidant enzyme genes of ROS modulation: gene name, product description and primer sequences, previously described by Peters et al. (2017).

### RNA extraction and RT-qPCR analysis

Total RNA was extracted using TRIzol™ Plus RNA Purification Kit (Thermo Fisher Scientific), following the manufacturer’s instructions. After extraction, total RNA was treated with DNase I (Sigma Aldrich), and RNA integrity was verified by agarose gel electrophoresis. The cDNA was prepared using the GoScript Reverse Transcription System (Promega) according to the manufacturer’s recommendations, using 1 μg of total RNA as input. RT-qPCR reactions were performed with the 7300 Fast Real-Time PCR System (Applied Biosystems) using the GoTaq® qPCR Master Mix Kit (Promega). All RT-qPCR reactions were performed using a primer concentration of 1 μM. Three biological replicates were evaluated, and the Mock (non-inoculated plants) were used for calibration in the analysis of plant genes. PCR efficiency and quantification cycle (Cq) values were calculated using the LinReg PCR program v.2017.0 [Ramakers et al., 2003]. The relative expression ratios were calculated by the Pfaffl method [Pfaffl, 2001], normalized according to Vandesompele et al. (2002) and Hellemans et al. (2007).

### Statistical analysis

Data normality was assessed using the Shapiro–Wilk test, and variance homogeneity using Levene’s test. Based on the results, Student’s t-test, Welch’s t-test, or the non-parametric Mann–Whitney U test was applied. Statistical significance was set at *p* < 0.05 (inoculated vs. mock, Vertix2 inoculated vs. Vertix1 inoculated or Time-point-specific vs. combined reference). All analyses were performed using the scipy.stats module from SciPy v.1.11.4.

### Analysis of reference gene expression stability

To identify the most stable reference genes, we evaluated gene stability using three complementary statistical approaches. First, we performed a direct assessment of raw Cq values by calculating their mean, standard deviation (SD), and coefficient of variation (CV), and ranked the genes according to their CV values. Second, we employed the RefFinder web-based tool (https://www.ciidirsinaloa.com.mx/RefFinder-master/) [Xie et al., 2023], which integrates four widely-used algorithms (geNorm, NormFinder, BestKeeper, and the comparative ΔCt method) to generate a comprehensive stability ranking. Third, we used the pairwise variation (V) values calculated by geNorm [Vandesompele et al., 2002] to determine the optimal number of reference genes required for robust data normalization across all samples and experimental conditions.

These analyses were performed independently on four datasets: samples collected at 12 hours post-inoculation (hpi), 48 hpi, and 5 days post-inoculation (dpi), as well as a combined set including all time points for each energy cane genotype (Table 3). To integrate these results into a consolidated stability hierarchy, we aggregated the rankings across all analyses using the RankAggreg R package v0.6.6 employing the Cross-Entropy Monte Carlo algorithm [Pihur et al., 2009].

**Table 3.**
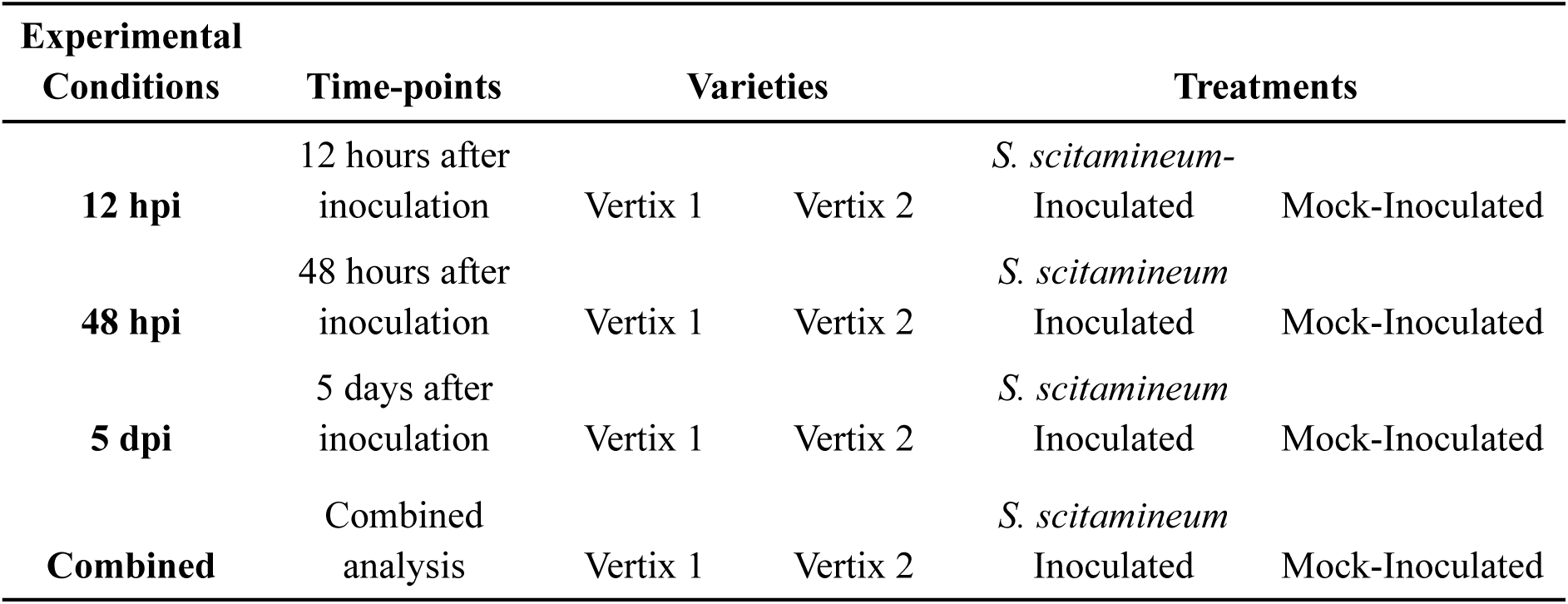
Summary of experimental conditions, detailing the energy cane varieties, treatments, and time-points used for the analysis.

## Results

### Selection of candidate reference genes using transcriptome data

Transcriptome data were analyzed to identify stably expressed genes across all 12 samples. Expression values were normalized by the TMM method, followed by transformation to counts per million (CPM), resulting in a dataset of 89,556 genes for the Inoculated Vx1 vs. Mock Vx1 (susceptible genotype) contrast, and 89,385 expressed genes for the Inoculated Vx2 *vs.* Mock Vx2 (resistant genotype) comparison (Suppl. File 1: Table S1). Principal component analysis (PCA) unveils a higher dispersion among Vx1 inoculated samples (PC1: 69%, PC2: 11%) and a clear separation between inoculated and Mock samples in Vx2 (PC1: 45%, PC2: 18%) (Suppl. File 2: Figure S1).

We identified a shared dataset of 82,848 genes (≈93% of all sugarcane genes) common to both genotypes (Suppl. File 1: Table S1). For each annotated gene, expression stability was inferred from the normalized counts of its aligned transcripts, and the coefficient of variation (CV) was used as a stability metric. Genes supported by transcript expression varying less than 10% among all twelve samples were ranked among the most stable, resulting in 2,147 shared genes for downstream analyses (Suppl. File 1: Table S2). Functional annotation based on the sugarcane R570 genome revealed two candidate references: SoffiXsponR570.05Eg030000 (AT5G66760; succinate dehydrogenase 1-1, CV 7.3%, mean 62.9 CPM) and SoffiXsponR570.5_9Ag072200 (AT1G13950; eukaryotic elongation factor 5A-1, CV 7.6%, mean 173.6 CPM). Considering possible allelic/paralogous copies in the polyploid sugarcane genome, BLASTn searches performed against the R570 genome, retrieved 10 and 4 homologs, respectively, that met the filtering criteria (expressed in both genotypes, CV ≤10%). These sequences were grouped into gene-specific sets (Suppl. File 1: Tables S3-S6), which were then used for primer design based on conserved regions identified through multiple sequence alignments (Suppl. File 2: Figures S2-S3)

### Validation of primer specificity and amplification efficiency

Primer specificity was confirmed using individual melting curves for each reference gene (Suppl. File 3: Figure S1), and quantification cycle (Cq) values were used for subsequent analyses (Suppl. File 4: Table S1). The regression coefficient (R^2^) for all primers varied between 0.997–1.000, and reaction efficiencies ranged from 83.9% to 100% for Vx1 (Suppl. File 4: Table S2), and 82.7% to 99.6% for Vx2 (Suppl. File 4: Table S3).

The expression levels of the nine candidate reference genes varied widely across both the Vx1 and Vx2 conditions, with mean Cq values ranging from 14.367 to 23.120 in Vx1 and from 14.529 to 23.327 in Vx2 (Figure 1). In Vx1, expression variability (ΔCq) was highest at 5 dpi (8.278) and 48 hpi (8.224), whereas for Vx2, the highest variability occurred at 5 dpi (7.534). In both experiments, the 12 hpi set showed the least variation (ΔCq = 5.688 and 5.652, respectively). Consistently across all samples, *SAMDC* was the most expressed gene (mean Cq = 14.367 ± 0.433 for Vx1; 14.529 ± 0.340 in Vx2), while *25S rRNA* was the least expressed (mean Cq = 23.120 ± 0.879 for Vx1; 23.327 ± 0.605 in Vx2).

**Fig. 1.**
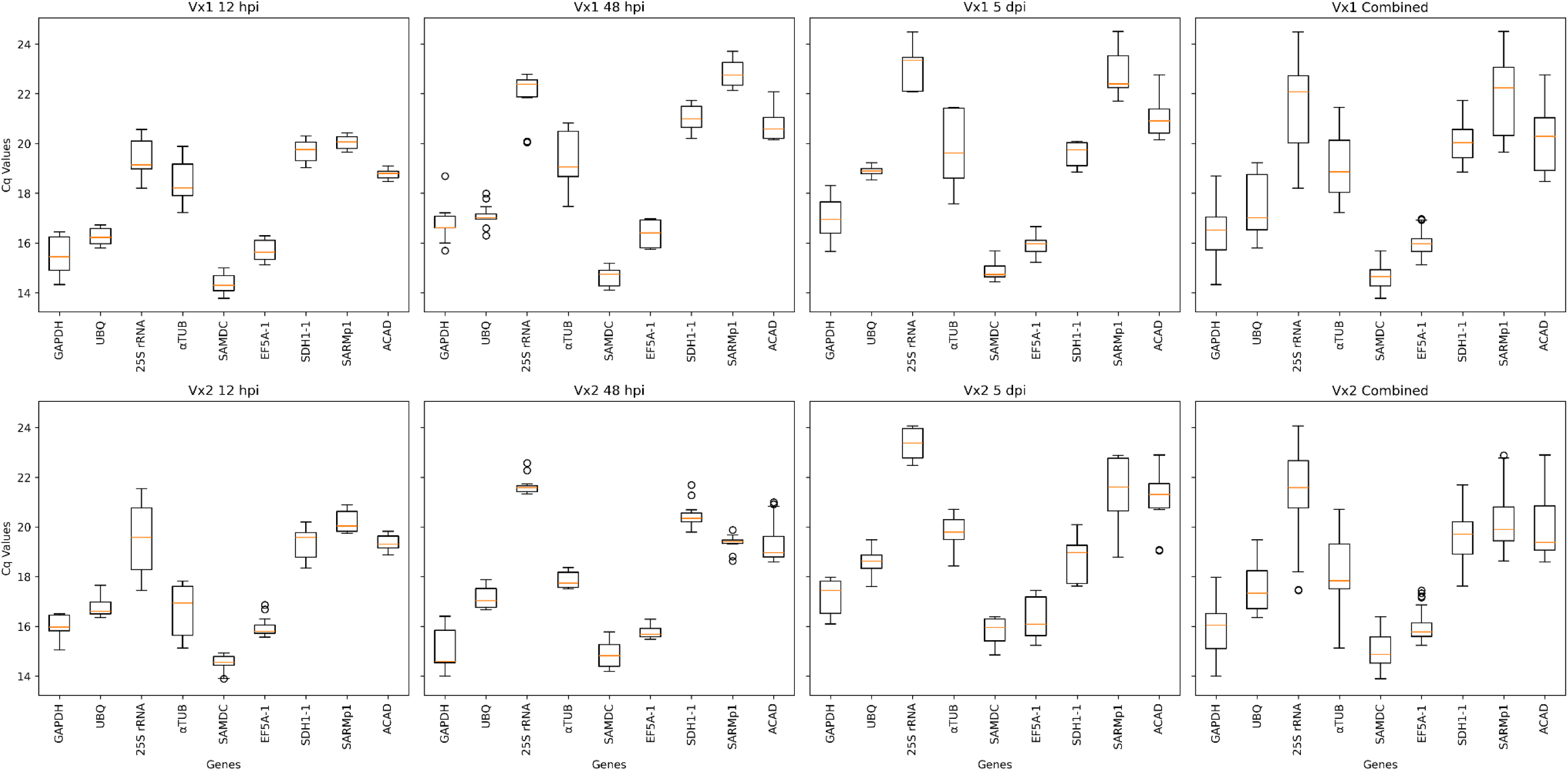
Distribution of Cq values for nine candidate reference genes across different time points and energy cane genotypes. Boxplots display the Cq value spread for each gene. The figure is organized into two rows by genotype (top: Vx1, bottom: Vx2) and four columns by treatment condition: 12 hpi, 48 hpi, 5 dpi, and Combined. Each box represents the interquartile range (IQR), with the horizontal line indicating the median Cq value and whiskers extending to 1.5× IQR

We assessed gene expression stability by calculating the coefficient of variation (CV) for each time point and for the combined dataset. Gene expression variability differed substantially across time points, conditions, and data subsets (Suppl. File 3: Figure S2). At 12 hpi, *ACAD* and *SARMp1* were the most stable genes in both Vx1 and Vx2, whereas *αTUB* and *25S rRNA* showed high variability. By 48 hpi, stability patterns diverged; in Vx1, a broad set of genes, including *25S rRNA* and *GAPDH,* were stable, while in Vx2, *EF5-1* and *SARMp1* were the most stable. At 5 dpi, *UBQ* showed high stability in both conditions. Finally, when analyzing the combined dataset, *SAMDC* and *EF5-1* were consistently the most stable genes for both Vx1 (CV = 3.07%–4.04%) and Vx2 (CV = 3.71%–4.84%), while *25S rRNA*, *SARMp1*, and *αTUB* were among the most variable (CV > 6.7%).

### RefFinder-based assessment of reference gene stability

Expression stability was individually evaluated using the RefFinder tool (Suppl. File 4: Table S4-S5). At 12 hpi, *ACAD* was consistently ranked as the most stable gene in both genotypes by most algorithms, followed by *SARMp1* in Vx1 and *SAMDC* in Vx2 (Figure 2). At 48 hpi, stability rankings diverged between genotypes: in Vx1, *SDH1-1*, *SAMDC*, and *EF5A-1* were identified as the most stable genes across all algorithms, whereas in Vx2, *EF5A-1*, *SARMp1*, and *αTUB* were top-ranked (Figure 2). At 5 dpi, *EF5A-1*, *UBQ*, and *SAMDC* were the most stable genes in Vx1, while *UBQ* and *25S rRNA* were the most stable in Vx2 (Figure 2). For the combined dataset including all time points, *SAMDC* remained the most stable gene in Vx1, whereas *UBQ* and *SAMDC* were consistently identified as the most stable genes in Vx2 (Figure 2)

**Fig. 2.**
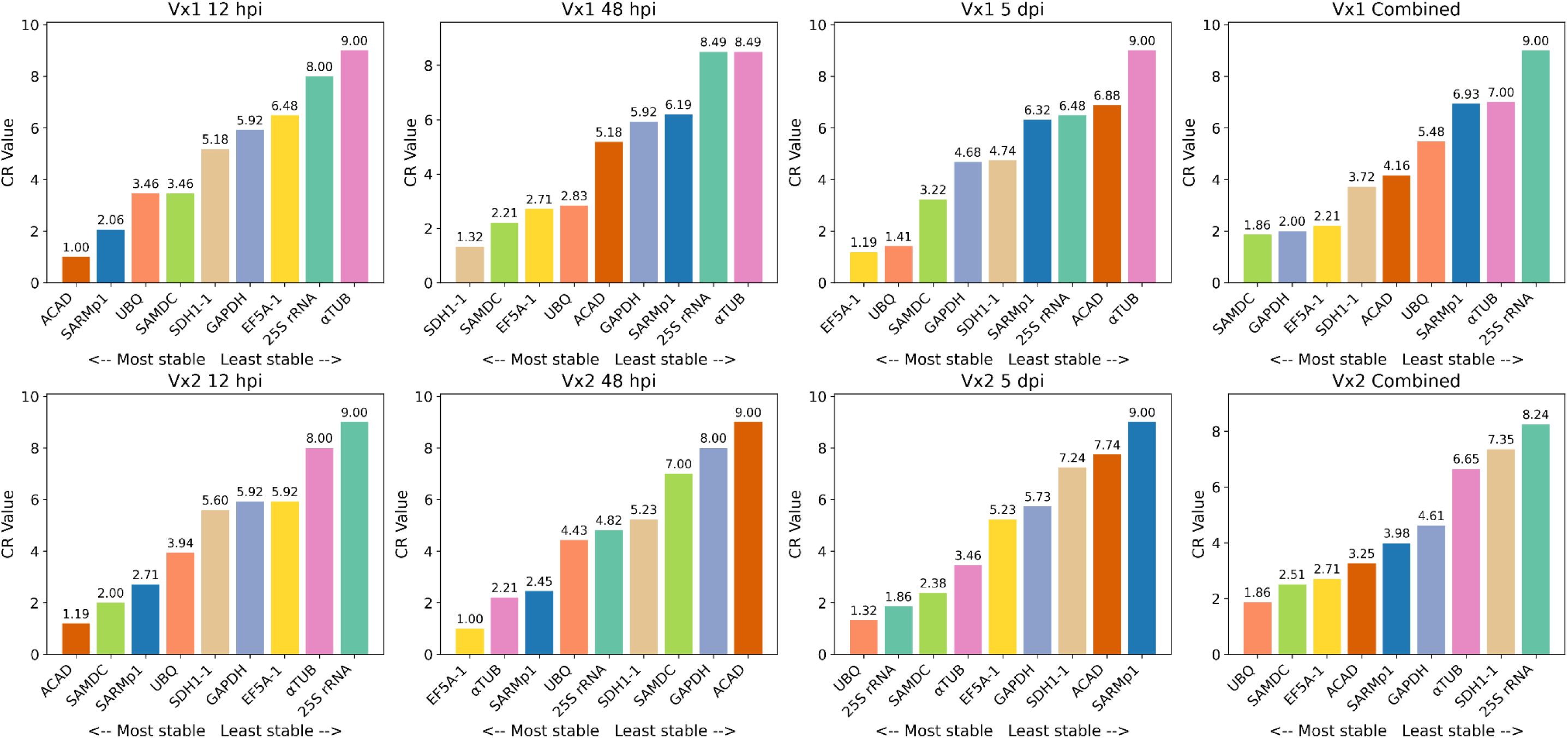
Stability ranking from RefFinder of candidate reference genes across different experimental conditions. Bar plots represent the geometric mean (Geomean) of comprehensive ranking (CR) values obtained from RefFinder analysis for nine candidate reference genes. Each subplot corresponds to a specific genotype (Vx1 or Vx2) and condition (12 hpi, 48 hpi, 5 dpi, and Combined). Lower CR values indicate higher expression stability. Bars are colored consistently across all subplots according to the gene identity: *GAPDH* (lavender), *UBQ* (light green), *25S rRNA* (light orange), *aTUB* (soft red), *SAMDC* (light gray-blue), *EF5-1* (light purple), and *SDH1-1* (pale blue). The gene order in each subplot is sorted from most stable (left) to least stable (right)

### Optimal combination of reference genes for expression normalization

We analyzed pairwise variation (Vn/Vn + 1) using geNorm to determine the optimal number of reference genes required for accurate normalization. All time-specific datasets (12 hpi, 48 hpi, and 5 dpi) for both genotypes showed pairwise variation values below the cutoff (< 0.15) [Vandesompele et al., 2002], indicating that the use of two reference genes is adequate for normalization (Figure 3). However, for the combined datasets, the minimum number of reference genes increased to three for Vx1 and four for Vx2.

**Fig. 3.**
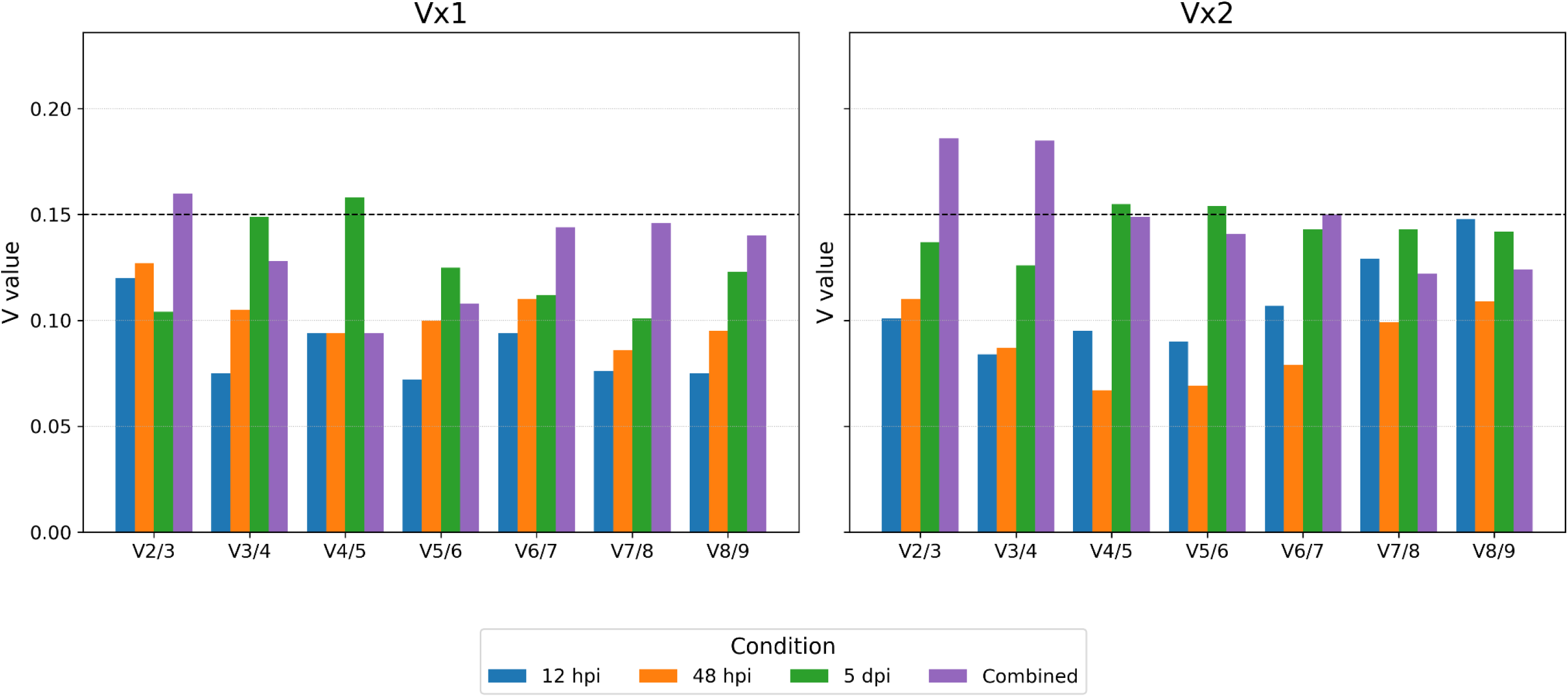
Pairwise variation (V) values across seven pairwise comparisons (V2/3 to V8/9) for two genotypes, Vx1 and Vx2. Bar plots show V values for four conditions measured at different time points post-inoculation: 12 hours post-inoculation (12 hpi, blue), 48 hours post-inoculation (48 hpi, orange), 5 days post-inoculation (5 dpi, green), and combined data (purple). The horizontal dashed line at V = 0.15 indicates the commonly used threshold for sufficient pairwise variation

The composition of optimal gene combinations varied depending on the time points analyzed (Suppl. Files 3: Figures S3–S4). At 12 hpi, *SARMp1* and *ACAD* were the most stable in both genotypes. At 48 hpi, *EF5-1* was common to both, with *αTUB* additionally stable in Vx2 and *SDH1-1* in Vx1. At 5 dpi, no overlap was observed: *25S rRNA* and *SAMDC* were the most stable in Vx2, whereas *UBQ* and *EF5-1* were recommended for Vx1. In the combined datasets, *EF5-1* and *SDH1-1* were consistently among the most stable across genotypes, with *25S rRNA* additionally selected for Vx1 and *UBQ* and *SAMDC* for Vx2.

### Consensus-based recommendation of reference genes

Considering all the metrics proposed (Lowest Coefficient of Variation, RefFinder, and geNorm), we consolidated our results into a final, condition-specific ranking presented in Table 4. The comprehensive RankAggreg output is available as supplementary file 4: Table S6. The optimal number of reference genes for each condition was determined by geNorm’s pairwise variation (V-value) analysis (Figure 3). At 12 hpi, a clear consensus emerged for both genotypes, identifying *ACAD* and *SARMp1* as the most stable candidates. In contrast, at 48 hpi, greater variation among methods was observed; *SDH1-1* and *SAMDC* were chosen as the final recommendations for Vx1, whereas *EF5-1* and *αTUB* were selected for Vx2. At 5 dai, *UBQ* consistently appeared among the top-ranked genes in both genotypes, paired with *EF5-1* for Vx1 and *25S rRNA* for Vx2. Finally, we observed a considerable overlap in stable genes across genotypes in the combined datasets. The candidates *SAMDC*, *EF5-1*, and *SDH1-1* were optimal for Vx1, and this same set was recommended for Vx2, with the addition of *UBQ*.

**Table 4.**
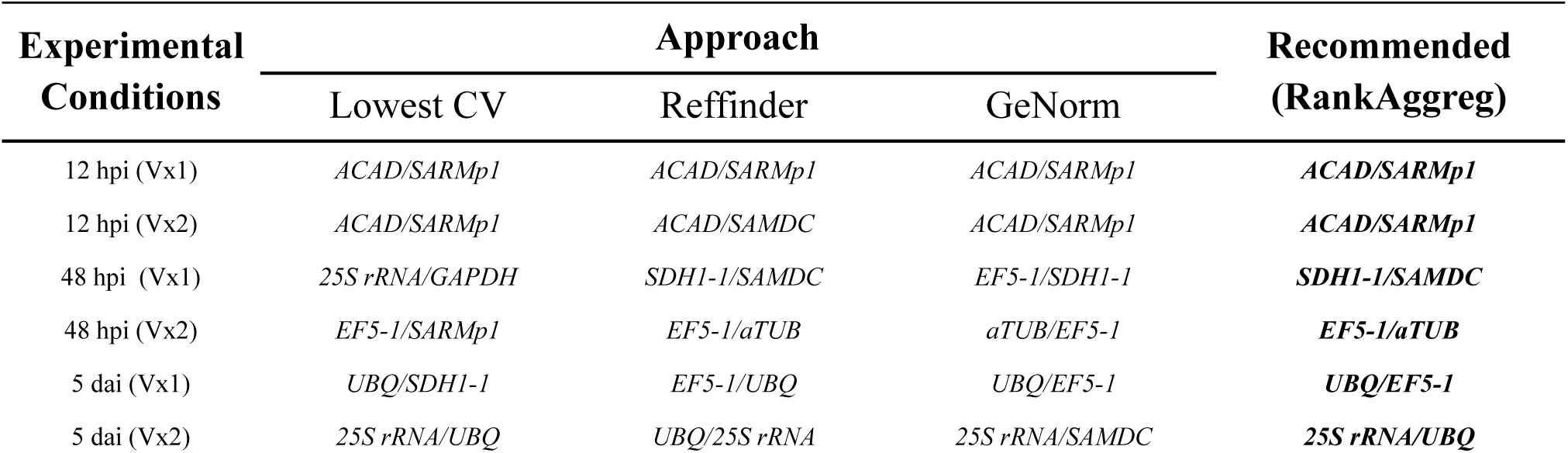
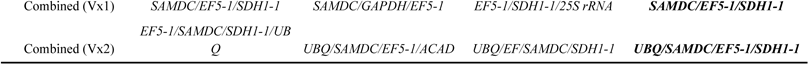
Final combinatorial recommendation of reference genes from RankAggreg for each experimental condition, according to three different approaches.

### Validation of reference genes using ROS-related gene expression analysis

To validate our optimal reference genes, we analyzed the expression patterns of a set of reactive oxygen species (ROS)-related defense genes: *SOD*, *CATB*, *POX5*, *GSTt3*, and *TRX*. Primer specificity was confirmed through the analysis of individual melting curves (Suppl. File 3: Figure S5). Gene expression data were normalized using either the specific set of recommended reference genes for each condition or the combined reference set determined independently of time point (Suppl. File 3: Figure S6).

The relative expression of the five ROS-related defense genes revealed clear time- and genotype-dependent transcriptional responses to *S. scitamineum* infection (Figure 4). In the resistant genotype Vx2, antioxidant enzymes, particularly *SOD* and *GSTt3*, were predominantly up-regulated across time points, while Vx1 exhibited delayed activation of these genes. *POX5* expression showed opposite regulation between genotypes, being mostly repressed in Vx1 but induced in Vx2 at 12 hpi and 5 dpi. *CATB* and *TRX* displayed moderate modulation, consistent with a coordinated antioxidant response in Vx2. Normalization using either the time-point-specific or combined reference gene sets generated comparable expression trends, confirming the robustness of the selected reference genes. Minor quantitative differences detected for *GSTt3* and *POX5* at 5 dpi in Vx2 (p < 0.05) did not alter the overall biological interpretation (Figure 4; Suppl. File 3: Figure S6; Suppl. File 4: Table S7).

**Fig. 4.**
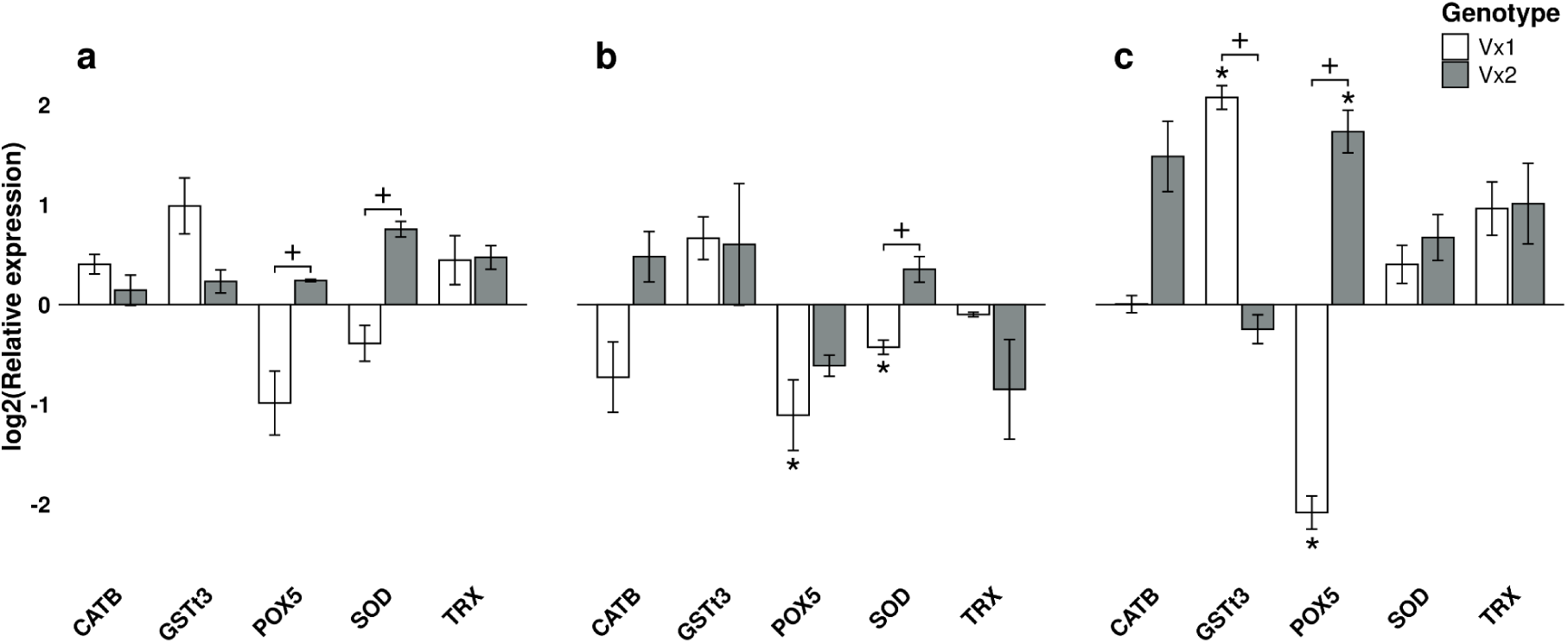
Expression profiles of five ROS-related genes in smut-susceptible and smut-resistant genotypes, measured by RT-qPCR at 12 hpi (a), 48 hpi (b), and 5 dpi (c) using time-point-specific reference genes. Asterisks (*) indicate genes that are significantly different between inoculated vs. mock. Plus signs (+) indicate genes that are significantly different between Vx2-inoculated vs. Vx1-inoculated plants. Error bars represent the standard error (SE)

Remarkably, when the same final combinatorial ranking approach was applied to identify the least stable reference genes(Suppl. File 4: Table S6), rather than the most stable ones, normalization of the ROS-related defense genes produced different outcomes. In the susceptible genotype Vx1, *SARMp1* and *aTUB* were selected for *POX5* and *SOD* at 48 hpi, and *GSTt3* at 5 dpi, whereas for Vx2, *SARMp1* and *ACAD* were selected for *POX5* at 5 dpi. When expression data were normalized using these genes, no ROS-related targets were detected as significantly different between inoculated vs. mock samples (Suppl. File 3: Figure S7).

## Discussion

This work provides the first systematic analysis of candidate reference genes for quantitative expression analyses in energy cane genotypes under *S. scitamineum* infection. We analyzed seven reference genes commonly used in various experiments, including those used in sugarcane-*S. scitamineum* interaction (*GAPDH, SAMDC, UBQ, 25S rRNA, αTUB, ACAD,* and *SARMp1),* together with two additional candidates (*EF5-1* and *SDH1-1*) identified from RNA-seq data of smut-infected energy cane genotypes. Using this transcriptomic dataset as a complementary screening step proved valuable for selecting genes suitable for energy cane expression studies.

Sugarcane and energy cane differ in genetic background and physiological traits [Diniz et al., 2019; Elumalai & Srinivasan, 2023; Hoang et al., 2015]. The two energy cane genotypes analyzed in this study, Vx1 and Vx2, have distinct parental lineages derived from different *Saccharum* spp. cultivars and *S. spontaneum* accessions, and exhibit genotype-dependent transcriptional responses to smut infection. This variability was consistent with our findings, as only the 12 hpi dataset shared a common set of reference genes between Vx1 and Vx2 (*ACAD* and *SARMp1*), indicating that gene expression stability in energy cane is genotype-dependent. Notably, these same genes were identified by Huang et al. (2018) as the most stable ones when comparing smut-infected sugarcane genotypes with contrasting resistance levels (NCo376 and YC71-374) at 3 and 7 dpi. Moreover, the commonly used sugarcane reference gene *GAPDH* exhibited considerable variation in expression levels between genotypes and across treatments. Although *GAPDH* is generally regarded as a reliable reference gene for universal sugarcane studies or those involving abiotic stress [Guo et al., 2014; Huang et al., 2018; Iskandar et al., 2004; Silva et al., 2014; Ling et al., 2014], its expression may fluctuate under pathogen infection [Huang et al., 2018]. Even though RNA-Seq data were available at a single time-point (48 hpi), *EF5-1* and *SDH1-1* were among the most stable genes across both genotypes and remained consistently stable at multiple time points [da Silva Santos et al., 2021; Marcolino-Gomes et al., 2015]. We identified endogenous genes that maintained stable expression across all treatments within each genotype, especially considering the marked distinct transcription profiles and variance patterns revealed by the PCA between resistant and susceptible genotypes. Using a similar approach, Huang et al. (2018) identified reliable reference genes for the sugarcane–smut interaction using expression microarray data, demonstrating that systematic validation of reference genes based on RNA-derived datasets is a valuable strategy for achieving accurate normalization in this pathosystem. Notably, when normalization was performed using the least stable candidates, several ROS-related genes lost statistical significance, leading to equivocal biological interpretations, showing that for sugarcane and energy cane, normalization gene selection is not a technical detail but a critical factor for detecting true transcriptional responses [Wang et al., 2021].

Reactive oxygen species are key mediators of plant immunity, acting as antimicrobial molecules and secondary messengers in defense signaling, activating several processes, such as strengthening the host cell wall via crosslinking glycoproteins [Mittler et al., 2017]. Upon inoculating *S. scitamineum*, resistant sugarcane plants trigger a localized oxidative burst, reflecting the early activation of basal plant defenses [Peters et al., 2017]. This timely ROS accumulation, together with the early induction of antioxidant enzymes, points to a tightly regulated balance between defense activation and cellular protection, one that appears compromised in the susceptible plants. To extend these observations, we investigated whether energy cane genotypes exhibiting contrasting responses to smut display similar transcriptional patterns in these early and late responsive ROS-associated defense genes.

Overall, the expression of antioxidant genes in energy cane showed a more variable pattern over time when compared with sugarcane. This divergence suggests that the timing of infection and the activation of defense responses may occur at different stages in the two pathosystems. The most considerable difference lies in the early and sustained induction of *SOD* in Vx2, whereas in Vx1, this gene remained repressed, in both cases, regardless of pathogen inoculation. SOD catalyzes the conversion of superoxide anion (O₂⁻•) into H₂O₂ and O₂, serving as the first line of defense against ROS [Ding et al., 2024; Lightfoot et al., 2017]. In contrast, in sugarcane, SOD expression and enzyme activity showed no significant variation between the resistant genotype SP80-3280 and the susceptible IAC66-6 after pathogen inoculation. However, SP80-3280 exhibits higher basal activity and a greater diversity of SOD isoenzymes than IAC66-6, independent of the inoculation or sampling time analyzed (6, 12, 24, 48, 72 hpi) [Peters et al., 2017]. Such temporal shifts could influence how each genotype coordinates its oxidative burst and detoxification mechanisms. A more detailed functional characterization, particularly assays of enzyme activity, as Peters et al. (2017) described, will be essential to clarify how these antioxidant systems contribute to disease resistance or susceptibility.

## Conclusion

In conclusion, this study provides the first rigorously validated set of reference genes for quantifying gene expression in energy cane during infection by *Sporisorium scitamineum*. Our work highlights the value of using RNA-Seq data to screen for novel candidates, a strategy that successfully identified highly stable genes and expanded the pool beyond those traditionally used. We demonstrated that overall reference gene stability is highly genotype- and time-dependent, with only the early 12 hpi time point sharing a common stable pair (*ACAD* and *SARMp1*) between the resistant (Vx2) and susceptible (Vx1) genotypes. Crucially, we show that the conventional reference gene *GAPDH* is unstable in this system, and its use can generate misleading interpretations of defense-related transcriptional changes. By applying our validated normalizers, we uncovered that the resistant genotype mounts a rapid and sustained antioxidant response (e.g., SOD induction) that is absent in the susceptible genotype. These findings provide an essential toolkit to ensure accuracy and reproducibility in future molecular studies, paving the way for a deeper understanding of the energy cane–smut pathosystem.

## Declarations

## Acknowledgements

We thank Elaine Vidotto Batista (Genomics Group Lab, ESALQ/USP) for technical support.

## Author information

### Authors and Affiliations

Department of Genetics, Luiz de Queiroz College of Agriculture, University of São Paulo, Piracicaba, São Paulo, Brazil

Gustavo Schiavone Crestana, Marcella Ferreira, Joyce Dellavechia Ferreti & Claudia Barros Monteiro-Vitorello

Sugarcane Center, IAC, Ribeirão Preto, São Paulo, Brazil Silvana Creste

José Bressiani

### Contributions

GSC, MF and JDF: Performed data analysis, conducted experiments, collected and interpretation of the data and manuscript writing, JAB and SC: Provided samples for validation experiments and manuscript reviewing and editing, CBMV: Conceived and designed the study and manuscript writing, reviewing and editing.

### Corresponding author

Correspondence to Claudia Barros Monteiro-Vitorello

### Ethics declarations

Ethics approval and consent to participate: Not applicable.

Consent for publications: Not applicable.

Competing interests: The authors declare that they have no competing interests.

