## Supplementary material for "Reference Gene Validation in Energy Cane Under Smut Infection and Insights into Antioxidant Gene Expression": Supplementary_File_2.pdf

a

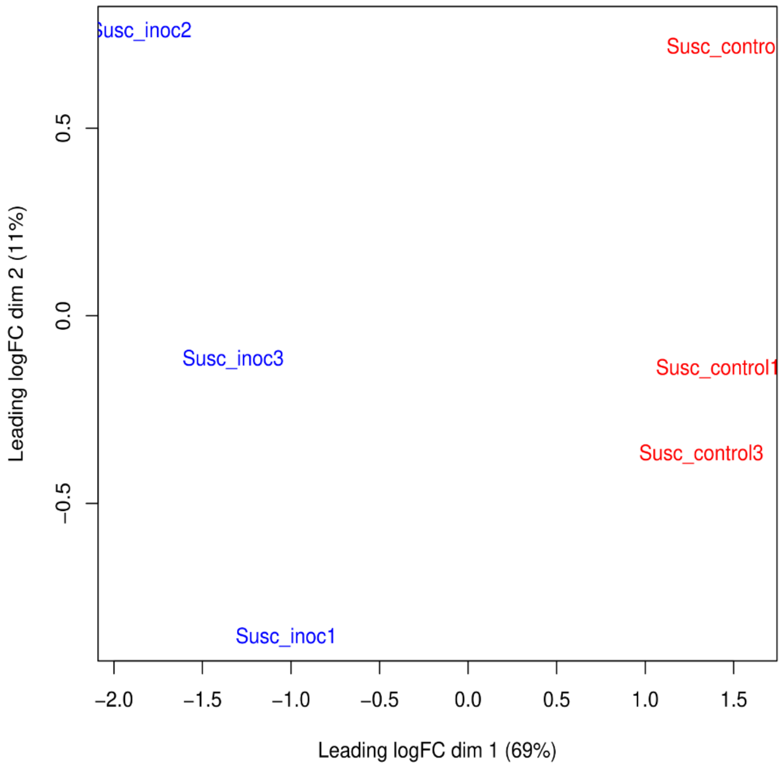


b

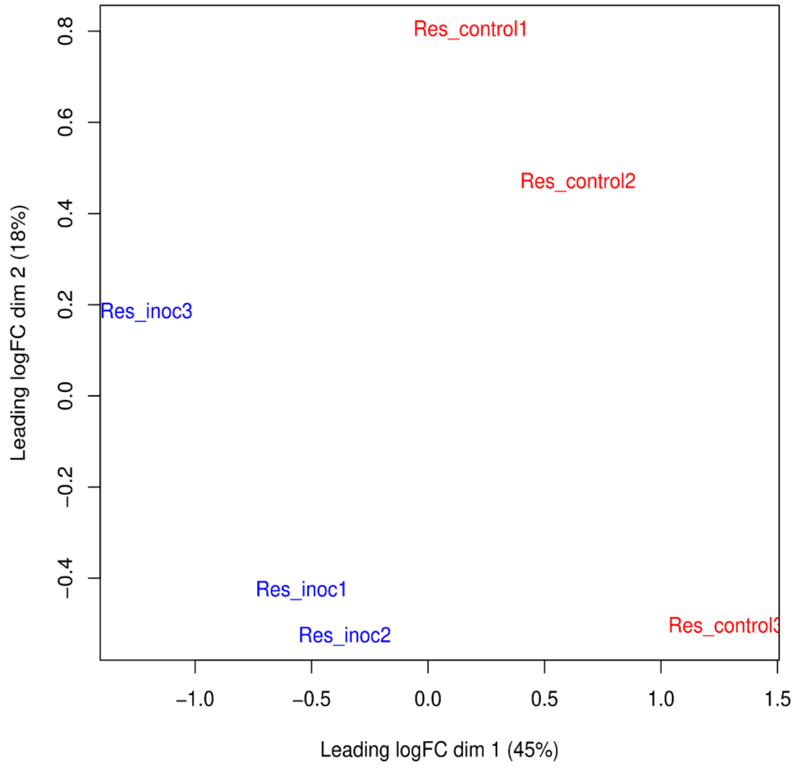

**Supplementary Fig. S1 Principal Component Analysis (PCA) of transcriptomic profiles from resistant and susceptible energy cane.** Clustering of control replicates (red) and replicates inoculated with *Sporisorium scitamineum* (blue) from the (A) susceptible genotype (Vx1) and the (B) resistant genotype (Vx2) . The percentage of total variance explained by each component is shown on the corresponding axis

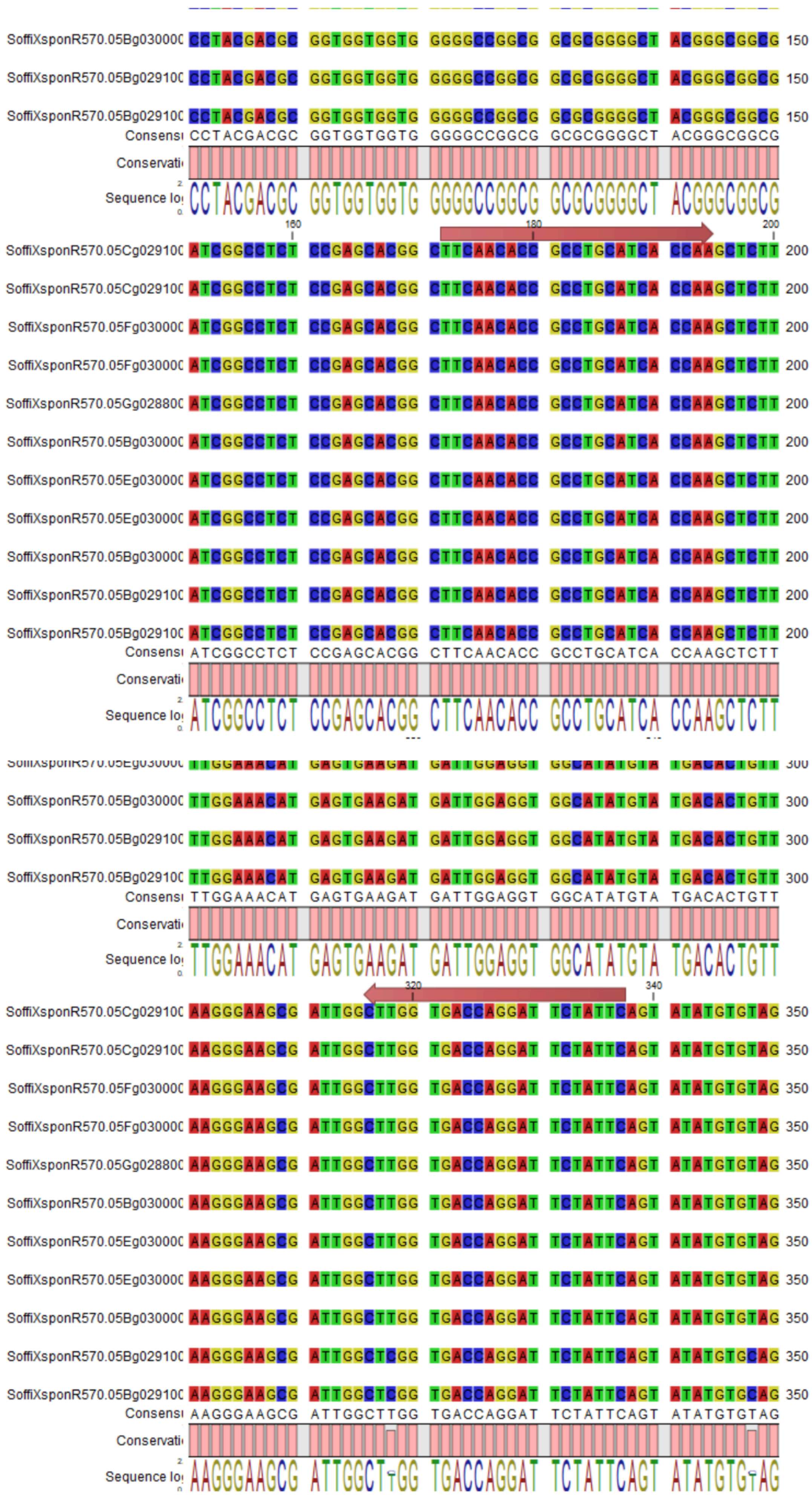

SoffiXsponR570.05Bg030000 CCTACGACGC GGTGGTGGTG GGGGCCGGCG GCGCGGGGCT ACGGGCGGCG 150
SoffiXsponR570.05Bg029100 CCTACGACGC GGTGGTGGTG GGGGCCGGCG GCGCGGGGCT ACGGGCGGCG 150
SoffiXsponR570.05Bg029100 CCTACGACGC GGTGGTGGTG GGGGCCGGCG GCGCGGGGCT ACGGGCGGCG 150
Consensus CCTACGACGC GGTGGTGGTG GGGGCCGGCG GCGCGGGGCT ACGGGCGGCG
Conservation
Sequence logo
160
180
200
SoffiXsponR570.05Cg029100 ATCGGCCTCT CCGAGCACGG CTTCAACACC GCCTGCATCA CCAAGCTCTT 200
SoffiXsponR570.05Cg029100 ATCGGCCTCT CCGAGCACGG CTTCAACACC GCCTGCATCA CCAAGCTCTT 200
SoffiXsponR570.05Fg030000 ATCGGCCTCT CCGAGCACGG CTTCAACACC GCCTGCATCA CCAAGCTCTT 200
SoffiXsponR570.05Fg030000 ATCGGCCTCT CCGAGCACGG CTTCAACACC GCCTGCATCA CCAAGCTCTT 200
SoffiXsponR570.05Gg028800 ATCGGCCTCT CCGAGCACGG CTTCAACACC GCCTGCATCA CCAAGCTCTT 200
SoffiXsponR570.05Bg030000 ATCGGCCTCT CCGAGCACGG CTTCAACACC GCCTGCATCA CCAAGCTCTT 200
SoffiXsponR570.05Eg030000 ATCGGCCTCT CCGAGCACGG CTTCAACACC GCCTGCATCA CCAAGCTCTT 200
SoffiXsponR570.05Eg030000 ATCGGCCTCT CCGAGCACGG CTTCAACACC GCCTGCATCA CCAAGCTCTT 200
SoffiXsponR570.05Bg030000 ATCGGCCTCT CCGAGCACGG CTTCAACACC GCCTGCATCA CCAAGCTCTT 200
SoffiXsponR570.05Bg029100 ATCGGCCTCT CCGAGCACGG CTTCAACACC GCCTGCATCA CCAAGCTCTT 200
SoffiXsponR570.05Bg029100 ATCGGCCTCT CCGAGCACGG CTTCAACACC GCCTGCATCA CCAAGCTCTT 200
Consensus ATCGGCCTCT CCGAGCACGG CTTCAACACC GCCTGCATCA CCAAGCTCTT
Conservation
Sequence logo
TTGGAAACAT GAGTGAAGAT GATTGGAGGT GGCATATGTA TGACACTGTT 300
SoffiXsponR570.05Bg030000 TTGGAAACAT GAGTGAAGAT GATTGGAGGT GGCATATGTA TGACACTGTT 300
SoffiXsponR570.05Bg029100 TTGGAAACAT GAGTGAAGAT GATTGGAGGT GGCATATGTA TGACACTGTT 300
SoffiXsponR570.05Bg029100 TTGGAAACAT GAGTGAAGAT GATTGGAGGT GGCATATGTA TGACACTGTT 300
Consensus TTGGAAACAT GAGTGAAGAT GATTGGAGGT GGCATATGTA TGACACTGTT
Conservation
Sequence logo
320
340
SoffiXsponR570.05Cg029100 AAGGGAAGCG ATTGGCTTGG TGACCAGGAT TCTATTCAGT ATATGTGTAG 350
SoffiXsponR570.05Cg029100 AAGGGAAGCG ATTGGCTTGG TGACCAGGAT TCTATTCAGT ATATGTGTAG 350
SoffiXsponR570.05Fg030000 AAGGGAAGCG ATTGGCTTGG TGACCAGGAT TCTATTCAGT ATATGTGTAG 350
SoffiXsponR570.05Fg030000 AAGGGAAGCG ATTGGCTTGG TGACCAGGAT TCTATTCAGT ATATGTGTAG 350
SoffiXsponR570.05Gg028800 AAGGGAAGCG ATTGGCTTGG TGACCAGGAT TCTATTCAGT ATATGTGTAG 350
SoffiXsponR570.05Bg030000 AAGGGAAGCG ATTGGCTTGG TGACCAGGAT TCTATTCAGT ATATGTGTAG 350
SoffiXsponR570.05Eg030000 AAGGGAAGCG ATTGGCTTGG TGACCAGGAT TCTATTCAGT ATATGTGTAG 350
SoffiXsponR570.05Eg030000 AAGGGAAGCG ATTGGCTTGG TGACCAGGAT TCTATTCAGT ATATGTGTAG 350
SoffiXsponR570.05Bg030000 AAGGGAAGCG ATTGGCTTGG TGACCAGGAT TCTATTCAGT ATATGTGTAG 350
SoffiXsponR570.05Bg029100 AAGGGAAGCG ATTGGCTCGG TGACCAGGAT TCTATTCAGT ATATGTGCAG 350
SoffiXsponR570.05Bg029100 AAGGGAAGCG ATTGGCTCGG TGACCAGGAT TCTATTCAGT ATATGTGCAG 350
Consensus AAGGGAAGCG ATTGGCTTGG TGACCAGGAT TCTATTCAGT ATATGTGTAG
Conservation
Sequence logo

**Supplementary Fig. S2 Multiple sequence alignment (MSA) for the SoffiXsponR570.05Eg030000 (AT5G66760) gene-specific dataset.** Nucleotide sequences (CDS) were aligned using ClustalW with CLC Genomics (default parameters), and predominantly conserved regions were used for primer design. Red annotation indicates the location of the forward (F) and reverse (R) primers within a conserved region

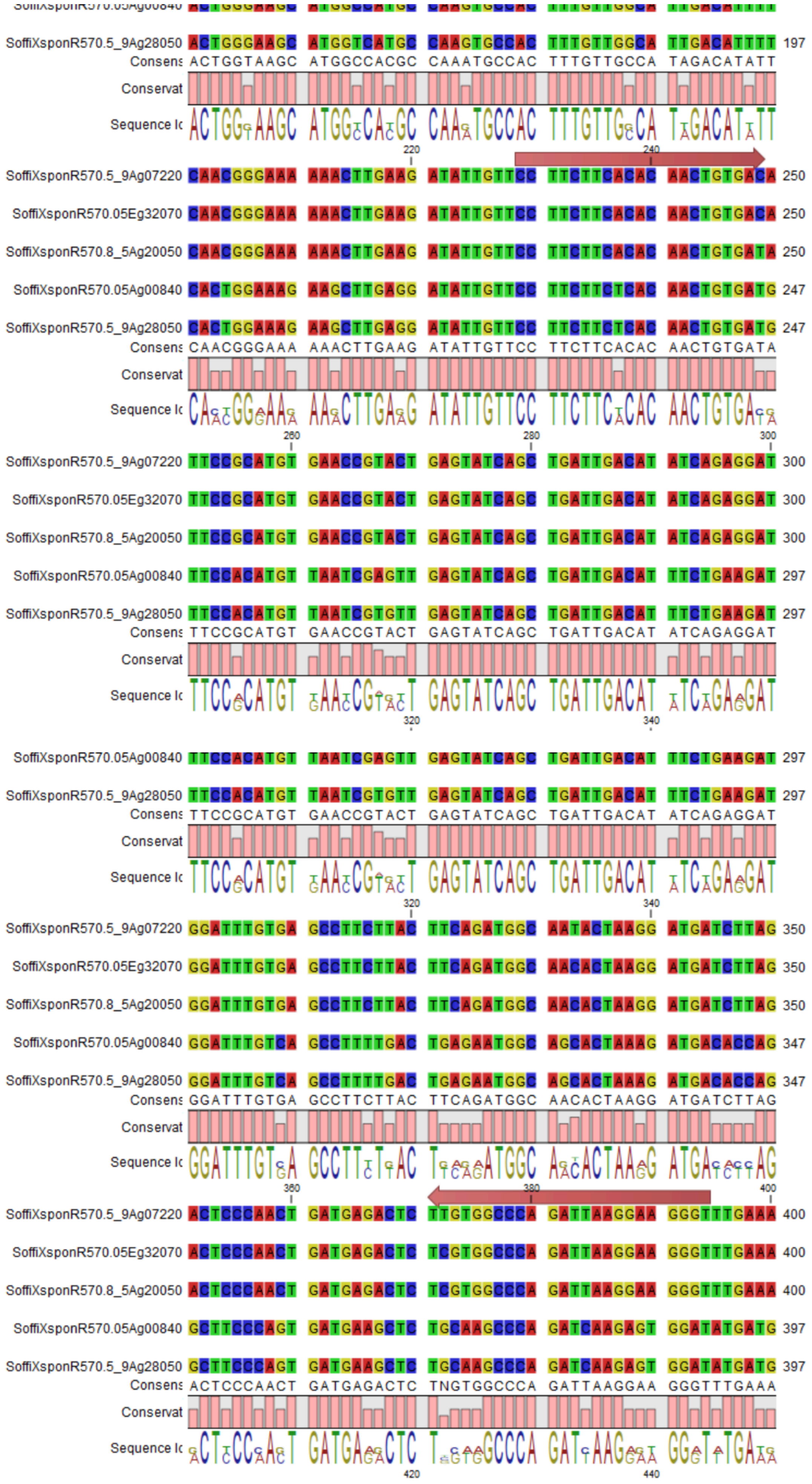
SoffiXsponR570.5_9Ag28050 ACTGGGAAGC ATGGTCATGC CAAGTGCCAC TTTGTTGGCA TTGACATTTT 197
Consens ACTGGTAAGC ATGGCCACGC CAAATGCCAC TTTGTTGCCA TAGACATATT
Conservat
Sequence lc
220 240
SoffiXsponR570.5_9Ag07220 CAACGGGAAA AAACTTGAAG ATATTGTTCC TTCTTCACAC AACTGTGACA 250
SoffiXsponR570.05Eg32070 CAACGGGAAA AAACTTGAAG ATATTGTTCC TTCTTCACAC AACTGTGACA 250
SoffiXsponR570.8_5Ag20050 CAACGGGAAA AAACTTGAAG ATATTGTTCC TTCTTCACAC AACTGTGATA 250
SoffiXsponR570.05Ag00840 CACTGGAAAG AAGCTTGAGG ATATTGTTCC TTCTTCTCAC AACTGTGATG 247
SoffiXsponR570.5_9Ag28050 CACTGGAAAG AAGCTTGAGG ATATTGTTCC TTCTTCTCAC AACTGTGATG 247
Consens CAACGGGAAA AAACTTGAAG ATATTGTTCC TTCTTCACAC AACTGTGATA
Conservat
Sequence lc
260 280 300
SoffiXsponR570.5_9Ag07220 TTCCGCATGT GAACCGTACT GAGTATCAGC TGATTGACAT ATCAGAGGAT 300
SoffiXsponR570.05Eg32070 TTCCGCATGT GAACCGTACT GAGTATCAGC TGATTGACAT ATCAGAGGAT 300
SoffiXsponR570.8_5Ag20050 TTCCGCATGT GAACCGTACT GAGTATCAGC TGATTGACAT ATCAGAGGAT 300
SoffiXsponR570.05Ag00840 TTCCACATGT TAATCGAGTT GAGTATCAGC TGATTGACAT TTCTGAAGAT 297
SoffiXsponR570.5_9Ag28050 TTCCACATGT TAATCGTGTT GAGTATCAGC TGATTGACAT TTCTGAAGAT 297
Consens TTCCGCATGT GAACCGTACT GAGTATCAGC TGATTGACAT ATCAGAGGAT
Conservat
Sequence lc
320 340
SoffiXsponR570.05Ag00840 TTCCACATGT TAATCGAGTT GAGTATCAGC TGATTGACAT TTCTGAAGAT 297
SoffiXsponR570.5_9Ag28050 TTCCACATGT TAATCGTGTT GAGTATCAGC TGATTGACAT TTCTGAAGAT 297
Consens TTCCGCATGT GAACCGTACT GAGTATCAGC TGATTGACAT ATCAGAGGAT
Conservat
Sequence lc
320 340
SoffiXsponR570.5_9Ag07220 GGATTTGTGA GCCTTCTTAC TTCAGATGGC AATACTAAGG ATGATCTTAG 350
SoffiXsponR570.05Eg32070 GGATTTGTGA GCCTTCTTAC TTCAGATGGC AACACTAAGG ATGATCTTAG 350
SoffiXsponR570.8_5Ag20050 GGATTTGTGA GCCTTCTTAC TTCAGATGGC AACACTAAGG ATGATCTTAG 350
SoffiXsponR570.05Ag00840 GGATTTGTCA GCCTTTTGAC TGAGAATGGC AGCACTAAAG ATGACACCAG 347
SoffiXsponR570.5_9Ag28050 GGATTTGTCA GCCTTTTGAC TGAGAATGGC AGCACTAAAG ATGACACCAG 347
Consens GGATTTGTGA GCCTTCTTAC TTCAGATGGC AACACTAAGG ATGATCTTAG
Conservat
Sequence lc
360 380 400
SoffiXsponR570.5_9Ag07220 ACTCCCAACT GATGAGACTC TTGTGGCCCA GATTAAGGAA GGGTTTGAAA 400
SoffiXsponR570.05Eg32070 ACTCCCAACT GATGAGACTC TCGTGGCCCA GATTAAGGAA GGGTTTGAAA 400
SoffiXsponR570.8_5Ag20050 ACTCCCAACT GATGAGACTC TCGTGGCCCA GATTAAGGAA GGGTTTGAAA 400
SoffiXsponR570.05Ag00840 GCTTCCCAGT GATGAAGCTC TGCAAGCCCA GATCAAGAGT GGATATGATG 397
SoffiXsponR570.5_9Ag28050 GCTTCCCAGT GATGAAGCTC TGCAAGCCCA GATCAAGAGT GGATATGATG 397
Consens ACTCCCAACT GATGAGACTC TNGTGGCCCA GATTAAGGAA GGGTTTGAAA
Conservat
Sequence lc
420 440

**Supplementary Fig. S3 Multiple sequence alignment (MSA) for the SoffiXsponR570.5_9Ag072200 (AT1G13950) gene-specific dataset.** Nucleotide sequences (CDS) were aligned using ClustalW with CLC Genomics (default parameters), and predominantly conserved regions were used for primer design. Red annotation indicates the location of the forward (F) and reverse (R) primers within a conserved region
