## Supplementary material for "Reference Gene Validation in Energy Cane Under Smut Infection and Insights into Antioxidant Gene Expression": Supplementary_File_3.pdf

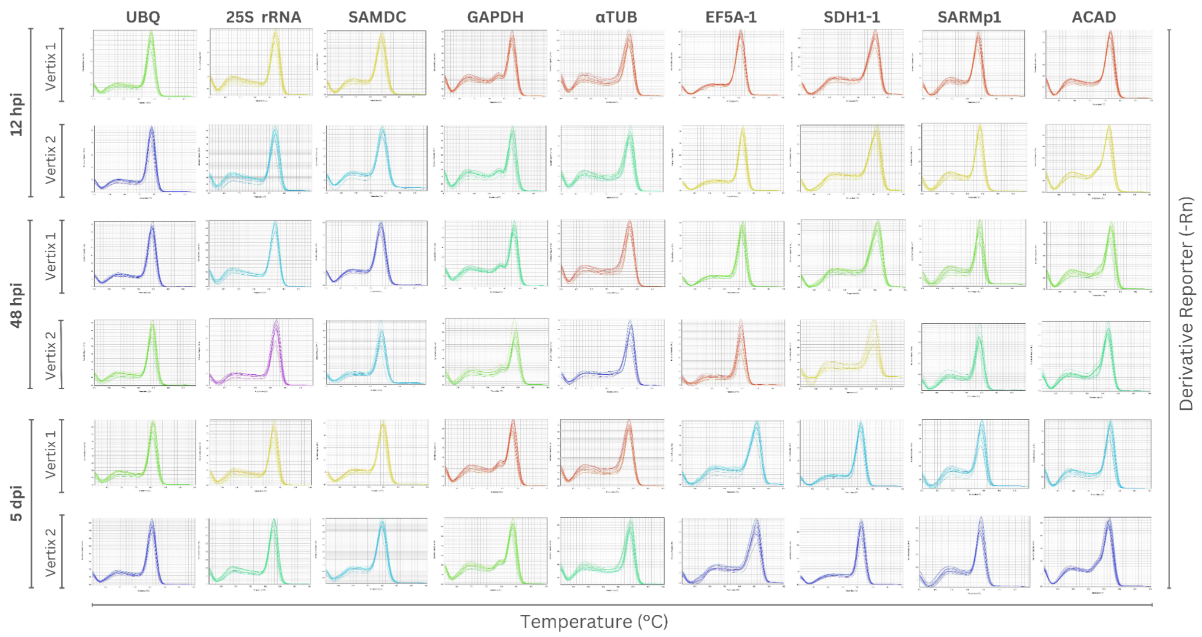


**Supplementary Fig. S1 Melting curve analysis confirming amplicon specificity for nine candidate reference genes.** Each curve represents a distinct melting peak across all time points for each gene, indicating specific amplification. The X-axis represents temperature (°C), and the Y-axis shows the derivative reporter signal (-Rn)

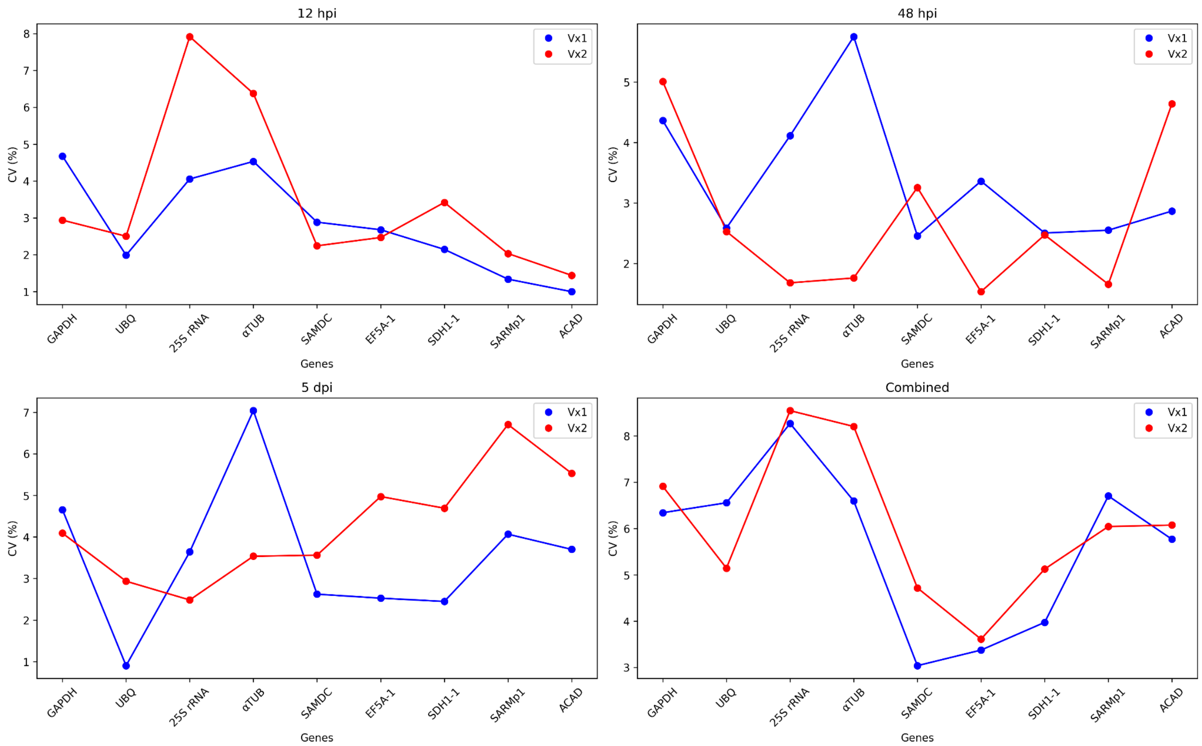


**Supplementary Fig. S2 Coefficient of variation (CV, %) of Cq values for nine candidate reference genes across time points in two energy cane genotypes.** CV values at 12 hours post-inoculation (hpi); 48 hpi; 5 days post-inoculation (dpi) and across all time points (Combined). Blue dots and lines represent genotype Vx1; red dots and lines represent genotype Vx2. Lower CV values indicate higher expression stability

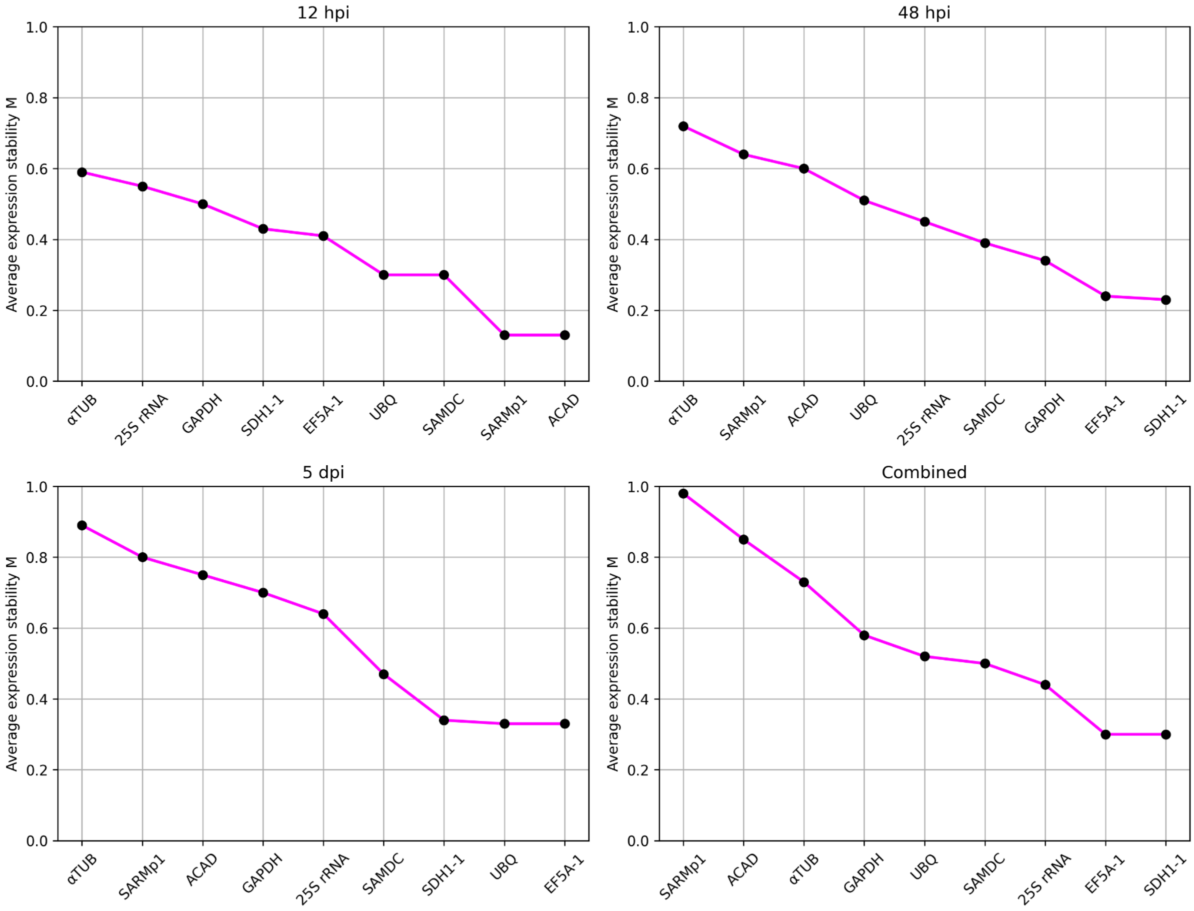


**Supplementary Fig. S3 Expression stability (M values) of candidate reference genes in genotype Vx1 across time points.** GeNorm analysis of nine candidate reference genes performed in Vx1 plants at different time points following pathogen inoculation: 12 hours post-inoculation (hpi), 48 hpi, 5 days post-inoculation (dpi), and a Combined dataset including all time points. Genes are ranked from least to most stable based on their average expression stability (M value), with lower M values indicating higher expression stability

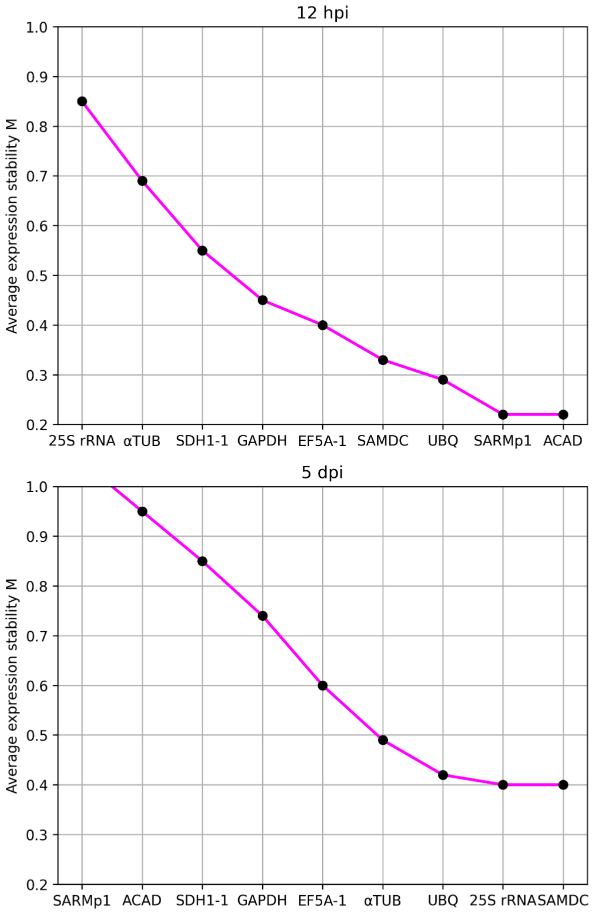


**Supplementary Fig. S4 Expression stability (M values) of candidate reference genes in genotype Vx2 across time points.** GeNorm analysis of nine candidate reference genes performed in Vx2 plants at different time points following pathogen inoculation: 12 hours post-inoculation (hpi), 48 hpi, 5 days post-inoculation (dpi), and a Combined dataset including all time points. Genes are ranked from least to most stable based on their average expression stability (M value), with lower M values indicating higher expression stability

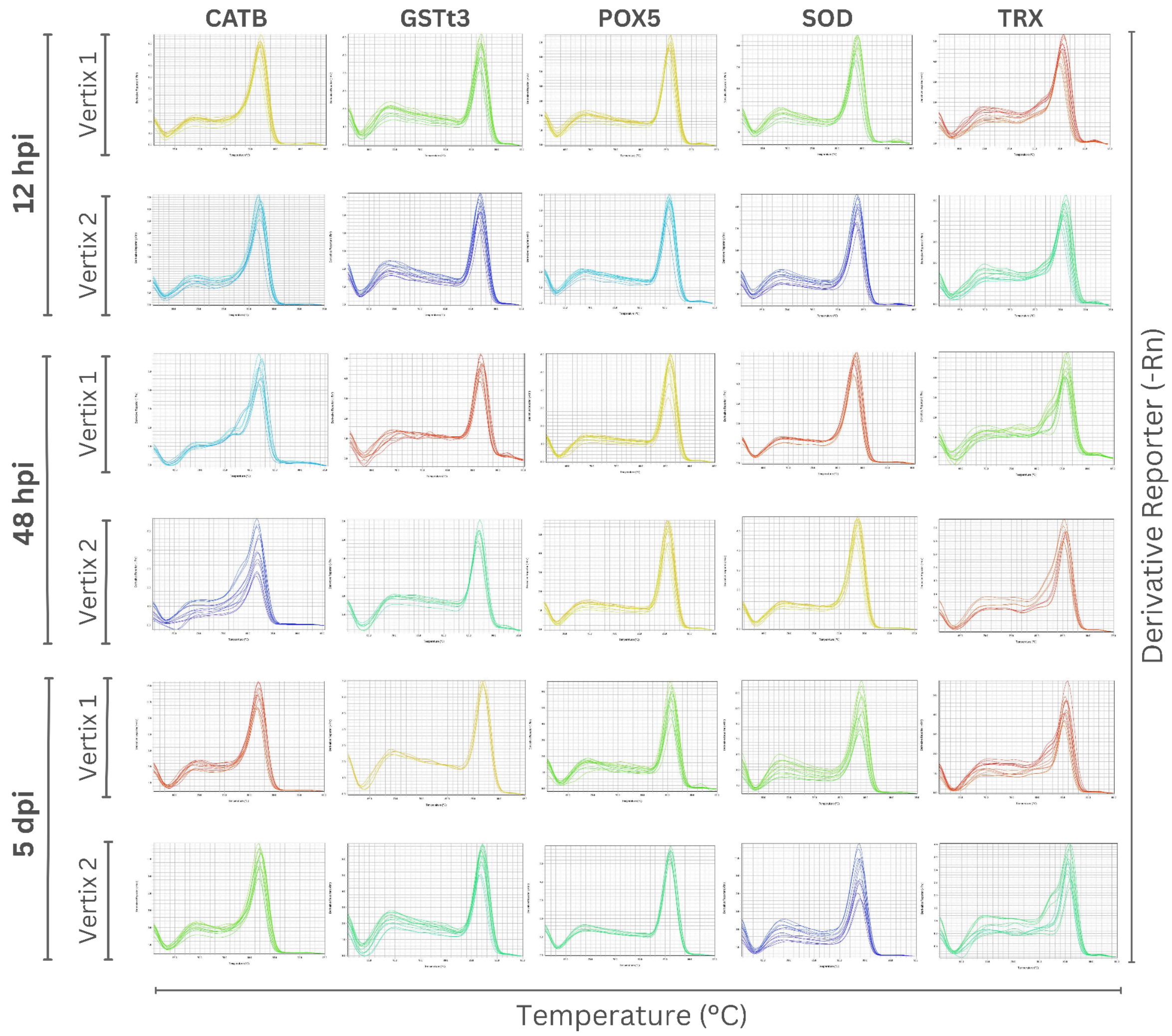


**Supplementary Fig. S5 Melting curve analysis confirming amplicon specificity for ROS metabolism genes**. Each curve represents a distinct melting peak across all time points for each gene, indicating specific amplification. The X-axis represents temperature (°C), and the Y-axis shows the derivative reporter signal (-Rn)

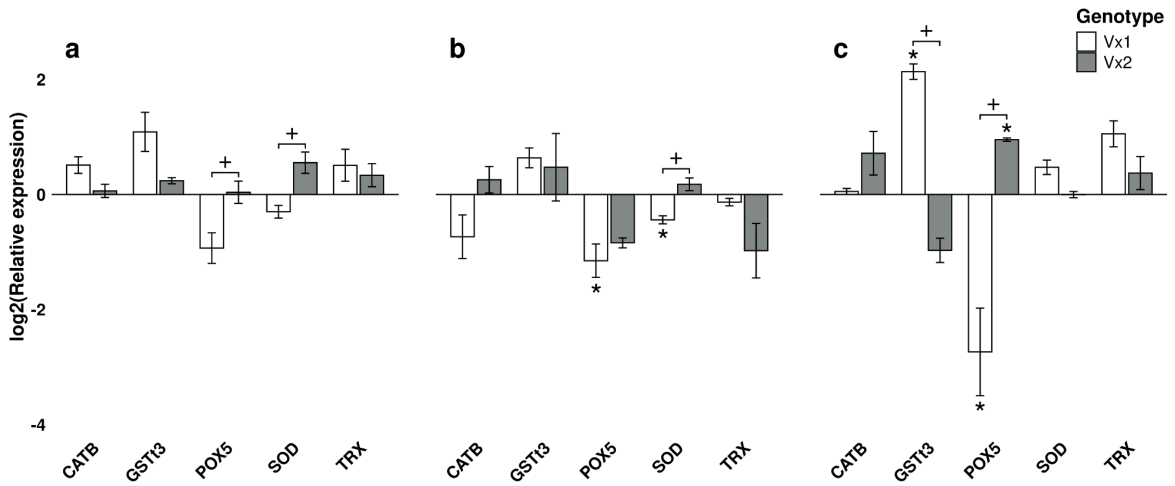


**Supplementary Fig. S6** Expression profiles of five ROS-related genes in smut-susceptible and smut-resistant genotypes, measured by RT-qPCR at 12 hpi (a), 48 hpi (b), and 5 dpi (c) using a combined reference gene set. Asterisks (*) indicate genes that are significantly different between inoculated vs. mock. Plus signs (+) indicate genes that are significantly different between Vx2-inoculated vs. Vx1-inoculated plants. Error bars represent the standard error (SE)

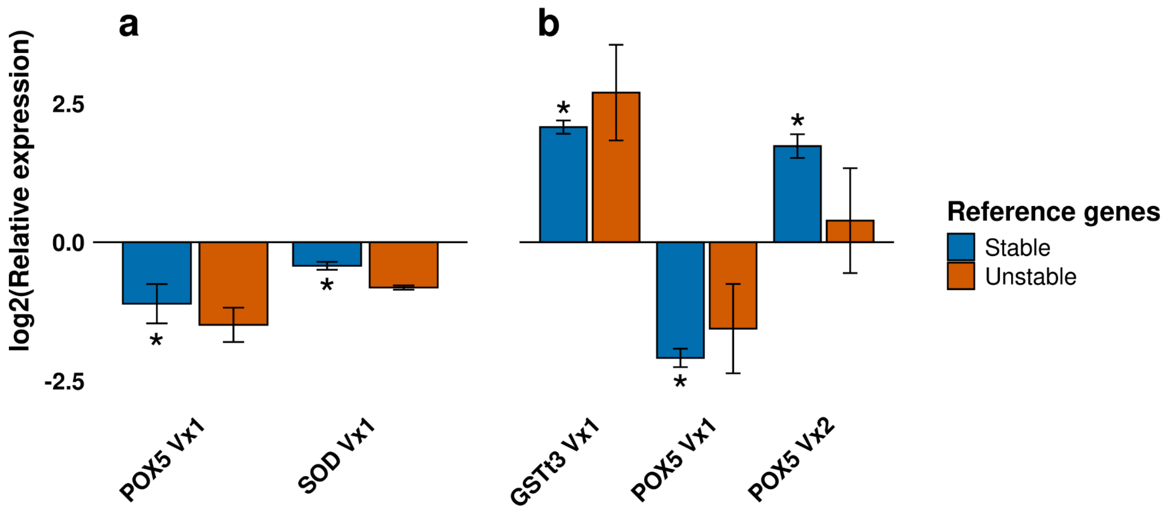


**Supplementary Fig. S7** Expression profile of ROS-related genes in smut-susceptible (Vx1) and -resistant (Vx2) genotypes by RT-qPCR analysis, normalized with stable vs. unstable reference genes. Gene expression at 48 hpi (a) and 5 dpi (b). Asterisks (*) indicate genes that are significantly different between inoculated vs. mock. Error bars represent the standard error (SE)
